# Cataloguing over-expressed genes in Epstein Barr Virus immortalized lymphoblastoid cell lines through consensus analysis of PacBio transcriptomes corroborates hypomethylation of chromosome 1

**DOI:** 10.1101/125823

**Authors:** Sandeep Chakraborty

## Abstract

The ability of Epstein Barr Virus (EBV) to transform resting cell B-cells into immortalized lymphoblastoid cell lines (LCL) provides a continuous source of peripheral blood lymphocytes that are used to model conditions in which these lymphocytes play a key role. Here, the PacBio generated transcriptome of three LCLs from a parent-daughter trio (SRAid:SRP036136) provided by a previous study [1] were analyzed using a kmer-based version of YeATS (KEATS). The set of over-expressed genes in these cell lines were determined based on a comparison with the PacBio transcriptome of twenty tissues provided by another study (hOPTRS) [2]. MIR155 long non-coding RNA (MIR155HG), Fc fragment of IgE receptor II (FCER2), T-cell leukemia/lymphoma 1A (TCL1A), and germinal center associated signaling and motility (GCSAM) were genes having the highest expression counts in the three LCLs with no expression in hOPTRS. Other over-expressed genes, having low expression in hOPTRS, were membrane spanning 4-domains A1 (MS4A1) and ribosomal protein S2 pseudogene 55 (RPS2P55). While some of these genes are known to be over-expressed in LCLs, this study provides a comprehensive cataloguing of such genes. A recent work involving a patient with EBV-positive large B-cell lymphoma was ‘unusually lacking various B-cell markers’, but over-expressing CD30 [3] - a gene ranked 79 among uniquely expressed genes here. Hypomethylation of chromosome 1 observed in EBV immortalized LCLs [4, 5] is also corroborated here by mapping the genes to chromosomes. Extending previous work identifying un-annotated genes [6], 80 genes were identified which are expressed in the three LCLs, not in hOPTRS, and missing in the GENCODE, RefSeq and RefSeqGene databases. KEATS introduces a method of determining expression counts based on a partitioning of the known annotated genes, has runtimes of a few hours on a personal workstation and provides detailed reports enabling proper debugging.

## Introduction

Epstein Barr Virus (EBV) transform resting cell B-cells into immortalized lymphoblastoid cell lines (LCL) [7], providing a continuous source of peripheral blood lymphocytes [8] to help model conditions in which these lymphocytes play a key role [9–11]. LCLs show high expression of several B-cell activation markers (FCER2, CD70, CD30, etc.) [12], and are extensively used to predict clinical response to anticancer drugs [13].

Pacific Biosciences (PacBio) sequencing [14] generates much longer reads compared to second-generation sequencing technologies [15, 16], with a trade-off of lower throughput, higher error rate and more cost per base [17, 18]. The longer sequence lengths in PacBio compared to other sequencing methods alleviate assembly issues associated with other methods with shorter read lengths [19, 20]. Unprecedented volumes of data generated by fast-evolving sequencing technologies necessitates the development of different pipelines to process and analyze this data. Transcriptomes are under-utilized while annotating genomes [21–23], as demonstrated on the walnut genome [24]. Previously, the MCF-7 transcriptome (2013 version, provided by Pacbio) was used to find transcripts that have no annotation in the current RefSeq and GENCODE databases, and predominantly absent in heart, liver and brain transcriptomes also provided by PacBio [6]. Also, shorter fragments of some of these transcripts were found to be present in seven tissues analyzed in a recent RACE-seq study (Accid:ERP012249) [25].

In the current work, three transcriptomes from a parent-daughter trio LCL cells lines (GM128LCLs) [1] were used to generate a consensus based catalogue of gene over-expressed in these cell lines as compared to the transcriptome from twenty different normal tissues (hOPTRS) [2]. This analysis required the development of an kmer-based assembly program within YeATS, named KEATS. KEATS identified several (n=765) genes that are expressed in GM128LCLs, but not found in hOPTRS. A recent work involving a patient with EBV-positive large B-cell lymphoma was ‘unusually lacking various B-cell markers’, but over-expressing CD30 [3] - a gene ranked 79 among uniquely expressed genes here. Furthermore, other genes (n=1361) were identified that had basal expression in hOPTRS, but higher expression in GM128LCLs. Hypomethylation of chromosome 1 observed in EBV immortalized LCLs [4, 5] is also corroborated here by mapping the genes to chromosomes. Extending previous work identifying un-annotated genes [6], 80 genes were identified which are expressed in the three LCLs, not in hOPTRS, and missing in the GENCODE, RefSeq and RefSeqGene databases. Thus, a catalogue of genes is generated that characterize LCLs, a model for studying many kinds of cancer.

## Results and discussion

Tilgner et al. [1] provided the PacBio transcriptome (SRAid:SRP036136) for three LCLs (GM128LCLs) from a parent-daughter trio (GM12878:n=715902, GM12891:n=586527 and GM12892:n=573590) [1], while another study has provided the PacBio transcriptome of a diverse pool of RNA samples representing 20 human tissues (hOPTRS) [2].

### Over-expressed genes in GM128LCLs with no expression in hOPTRS

Table 1 enumerates the first twenty genes with no corresponding transcripts in hOPTRS (see FILE:overExpressedCutoff10.txt for the complete list, n=765). The MIR155 gene, encoding the MiR-155 microRNA and the largest overexpressed gene in the GM128LCLs, is a widely studied gene known to promote the development and aggressiveness of B cell malignancies [26–29]. Another study using Northern blotting demonstrated that MIR155 has a 10 to 30 fold higher copy number in LCLs than in normal circulating B cells [30].

**Table 1:** Transcripts found in all three cell lines (GM12878, GM12891 and GM12892), and not hOPTRS. Transcripts are sorted based on cumulative counts. There are no corresponding transcripts in hOPTRS (see FILE:overExpressedCutoff0.txt for the complete list, n=765). Another method uses a count cutoff of 10 (Table 2). Cnt1: count in GM12878, Cnt2: count in GM12891, Cnt3: count in GM12892, Cumu: cumulative counts.

| Rank | Accid | Len | Description | Cnt1 | Cnt2 | Cnt3 | Cumu |
| --- | --- | --- | --- | --- | --- | --- | --- |
| 1 | NR_001458.3 | 1500 | MIR155 host gene (MIR155HG), lncRNA | 280 | 284 | 385 | 949 |
| 2 | NM_001207019.2 | 1488 | Fc fragment of IgE receptor II (FCER2), | 333 | 279 | 255 | 867 |
| 3 | NM_001098725.1 | 1405 | T-cell leukemia/lymphoma 1A (TCL1A), | 77 | 43 | 20 | 140 |
| 4 | NM_001190259.1 | 3355 | germinal center associated signaling and motility (GCSAM), | 34 | 30 | 50 | 114 |
| 5 | NM_016095.2 | 1196 | GINS complex subunit 2 (GINS2), | 34 | 31 | 48 | 113 |
| 6 | NM_001330332.1 | 2431 | CD70 molecule (CD70), | 40 | 41 | 29 | 110 |
| 7 | NM_001303618.1 | 2842 | CD226 molecule (CD226), | 33 | 25 | 37 | 95 |
| 8 | NM_005098.3 | 2075 | musculin (MSC), | 35 | 27 | 28 | 90 |
| 9 | NM_001142703.1 | 3407 | family with sequence similarity 111 member B (FAM111B), | 26 | 19 | 41 | 86 |
| 10 | NM_002915.3 | 2396 | replication factor C subunit 3 (RFC3), | 30 | 31 | 23 | 84 |
| 11 | NM_001284244.1 | 1022 | prepronociceptin (PNOC), | 41 | 28 | 15 | 84 |
| 12 | NR_046713.1 | 807 | NAALADL2 antisense RNA 2 (NAALADL2-AS2), lncRNA | 5 | 26 | 52 | 83 |
| 13 | NM_001146015.1 | 3084 | DLG associated protein 5 (DLGAP5), | 31 | 24 | 28 | 83 |
| 14 | NM_001267038.1 | 1719 | centromere protein K (CENPK), | 22 | 31 | 30 | 83 |
| 15 | NM_000594.3 | 1686 | tumor necrosis factor (TNF), | 23 | 32 | 25 | 80 |
| 16 | NM_001178098.1 | 1968 | CD19 molecule (CD19), | 31 | 31 | 14 | 76 |
| 17 | NM_001085357.1 | 3072 | B and T lymphocyte associated (BTLA), | 28 | 20 | 19 | 67 |
| 18 | NM_001286449.1 | 1782 | testis expressed 9 (TEX9), | 23 | 8 | 35 | 66 |
| 19 | NM_001306151.1 | 3006 | adaptor of phosphotyrosine and 3-phosphoinositides 1 (DAPP1), | 25 | 22 | 18 | 65 |
| 20 | XM_005269743.3 | 1491 | PREDICTED: H6 family homeobox 2 (HMX2), X1, | 11 | 30 | 24 | 65 |
| 79 | NM_001243.4 | 3706 | TNF receptor superfamily member 8 (TNFRSF8), | 11 | 4 | 11 | 26 |

### Over-expressed genes in GM128LCLs with basal expression in hOPTRS

Table 2 shows transcripts wherein the counts in hOPTRS are <10, and counts in GM128LCLs >10 (see FILE:overExpressedCutoff10.txt for the complete list, n=1361). 10 is used as an empirical cutoff. Ideally a statistic should be used to check for over-expressed genes, but will not significantly alter the rankings of the top-ranked genes presented here. MS4A1, the most over-expressed gene, encodes the B-lymphocyte antigen CD20 expressed ubiquitously on the surface B-cells in almost all stages. Anti-CD20 monoclonal antibodies are used for the treatment of patients with B-Cell malignancies [31], although CD20 was shown to have no prognostic value in acute lymphoblastoid leukemia [32].

**Table 2:** Transcripts found in all three cell lines (GM12878, GM12891 and GM12892) with count >10, and count <10 in hOPTRS. Transcripts are sorted based on cumulative counts. Here, the corresponding transcript counts in the hOPTRS are <10 (see FILE:overExpressedCutoff10.txt for the complete list, n=1361). Accid:NG_011221.1 marked with a single asterisk is annotated in RefSeq under a different ‘facet’, and thus not downloaded automatically. Accid:AL121985.13 marked with a double asterisk is not annotated in RefSeq, but is annotated in GENCODE (Id:OTTHUMT00000479908). Cnt1: count in GM12878, Cnt2: count in GM12891, Cnt3: count in GM12892, Cumu: cumulative counts.

| Rank | Accid | Length | Description | Cnt1 | Cnt2 | Cnt3 | Cumu |
| --- | --- | --- | --- | --- | --- | --- | --- |
| 1 | NR_001458.3 | 1500 | MIR155 host gene (MIR155HG), lncRNA | 280 | 284 | 385 | 949 |
| 2 | NM_001207019.2 | 1488 | Fc fragment of IgE receptor II (FCER2), mRNA | 333 | 279 | 255 | 867 |
| 3 | NM_152866.2 | 3594 | membrane spanning 4-domains A1 (MS4A1), mRNA | 225 | 210 | 400 | 835 |
| 4 | *NG_011221.1 | 883 | ribosomal protein S2 pseudogene 55 (RPS2P55) | 384 | 354 | 41 | 779 |
| 5 | NM_005101.3 | 685 | ISG15 ubiquitin-like modifier (ISG15), mRNA | 190 | 207 | 152 | 549 |
| 6 | NM_001256030.1 | 1537 | CD48 molecule (CD48), mRNA | 200 | 159 | 181 | 540 |
| 7 | NR_003098.1 | 1134 | small nucleolar RNA host gene 1 (SNHG1), lncRNA | 151 | 124 | 182 | 457 |
| 8 | NM_016459.3 | 845 | marginal zone B and B1 cell specific protein (MZB1), mRNA | 198 | 134 | 119 | 451 |
| 9 | NM_014508.2 | 1127 | apolipoprotein B mRNA editing enzyme (APOBEC3C), mRNA | 159 | 146 | 112 | 417 |
| 10 | NM_001143995.2 | 1965 | leupaxin (LPXN), mRNA | 124 | 112 | 178 | 414 |
| 11 | **AL121985.13 | 1370 | sequence from clone RP11-404F10 | 209 | 164 | 157 | 530 |
| 12 | NM_006417.4 | 1742 | interferon induced protein 44 (IFI44), mRNA | 110 | 119 | 121 | 350 |
| 13 | NM_005782.3 | 1113 | Aly/REF export factor (ALYREF), mRNA | 127 | 106 | 98 | 331 |
| 14 | NM_000265.5 | 1475 | neutrophil cytosolic factor 1 (NCF1), mRNA | 129 | 101 | 96 | 326 |
| 15 | NM_001184866.1 | 2375 | Fc receptor like A (FCRLA), mRNA | 144 | 89 | 83 | 316 |
| 16 | NM_001114735.1 | 955 | BCL2 related protein A1 (BCL2A1), mRNA | 78 | 84 | 150 | 312 |
| 17 | NM_002664.2 | 2866 | pleckstrin (PLEK), mRNA | 115 | 117 | 75 | 307 |
| 18 | NM_002358.3 | 1453 | MAD2 mitotic arrest deficient-like 1 (yeast) (MAD2L1), mRNA | 102 | 96 | 96 | 294 |
| 19 | NM_001039517.1 | 1326 | RUSC1 antisense RNA 1 (RUSC1-AS1), mRNA | 93 | 96 | 92 | 281 |
| 20 | NM_004418.3 | 1708 | dual specificity phosphatase 2 (DUSP2), mRNA | 74 | 74 | 130 | 278 |

### Genes assigned to chromosome corroborates the hypomethylation of chromosome 1

Table 3 shows that chromosome 1, known to be hypomethylated in EBV immortalized LCLs [4, 5], overexpressed the maximum number of genes. It has been shown that demethylation of satellite 3 DNA in chromosome 1 leads to increased transcription in senescent cells and in A431 epithelial carcinoma cells [33]. Hypomethylation of chromosome 1 and 16 have also been linked to Wilms tumors [34]. Also, ‘chromosome 1 is involved in quantitative anomalies in 50-60% of breast tumours’ [35], with three common genes (MLLT11, MTX1 and HIV-1) from chromosome 1 being reported here as being overexpressed (see FILE:overExpressedCutoff10.txt).

**Table 3:**
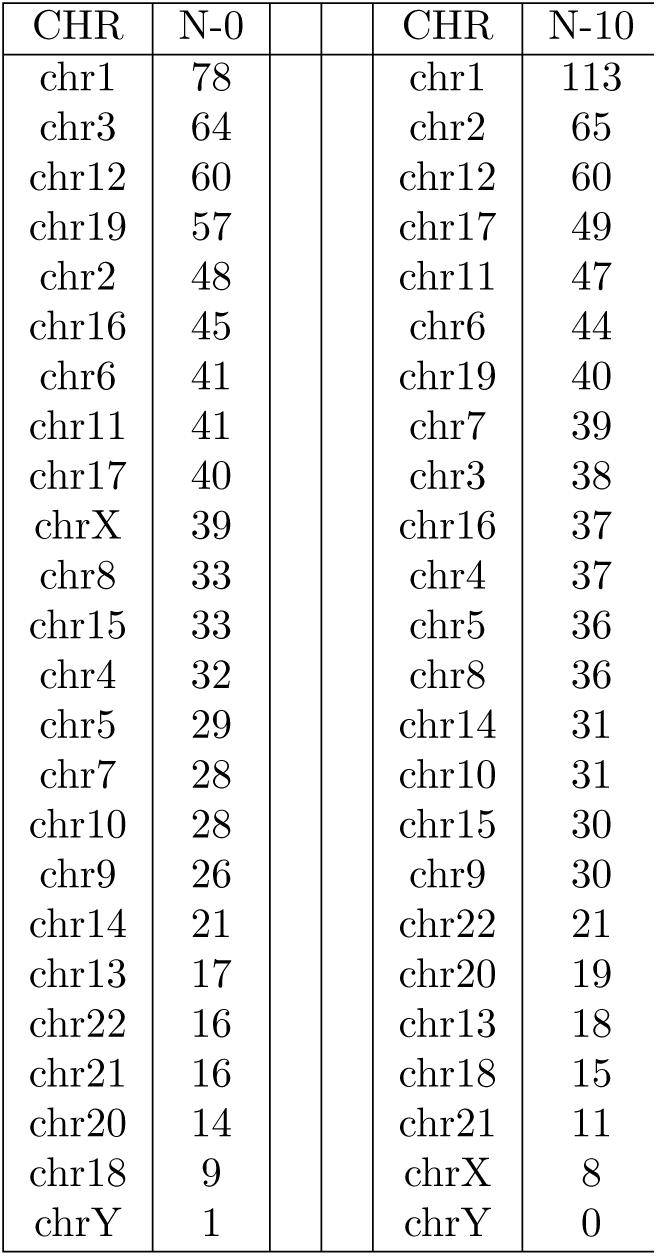
Over-expressed genes assigned to chromosomes. Chromosomes are sorted based on the number of over-expressed genes for a cutoff of 0 (N-0) and 10 (N-10). Chromosome 1 is known to be hypomethylated in EBV immortalized LCLs [4, 5], linked to Wilms tumors [34] and has increased transcription in senescent cells and in A431 epithelial carcinoma cells [33].

### Issues with RefSeq

In Table 2, Accid:NG_011221.1 marked with a single asterisk is annotated in RefSeq under a different ‘facet’, and thus not downloaded automatically. Ideally, this should have been part of the ‘mRNA’ facet. This can be an issue while benchmarking RefSeq with GENCODE [36]. Accid:AL121985.13 marked with a double asterisk is not annotated in RefSeq, but is annotated in GENCODE (Id:OTTHUMT00000479908). This gene is antisense to the CD48 antigen, a protein found on the surface of immune cells [37].

### The utility of a comprehensive catalogue

A recent work involving EBV-positive large B-cell lymphoma (DLBCL) was found to be lacking various standard B-cell markers, but over-expressing CD30/TNFRSF8 [3]. This study identified that the DLBCL case was ‘positive for CD30 and MUM-1, not defining the lineage of tumor cells’ [3]. However, a previous study had reported ‘CD30 was expressed in 14% of DLBCL patients. Patients with CD30+ DLBCL had superior 5-year overall survival’ [38]. Irrespective, the current study identifies CD30/TNFRSF8 as a gene uniquely expressed in LCLs, and its ranking shows that there are at least 78 other possible biomarkers (although many of them, like MIR155, are known and well established).

### Expression counts - detailed reporting in KEATS

KEATS provides a detailed reporting system to enable debugging results. Take NM_001243.4 (length=3706) in Table 1 - this has a 24 count in GM12878. The transcripts matching these are reported in a file which includes the lengths of the individual transcripts (Table 4). Normalization is achieved by dividing the sum of the lengths of these by the length of NM_001243.4, leading to count of 11 in GM12878 (Table 1).

**Table 4:**
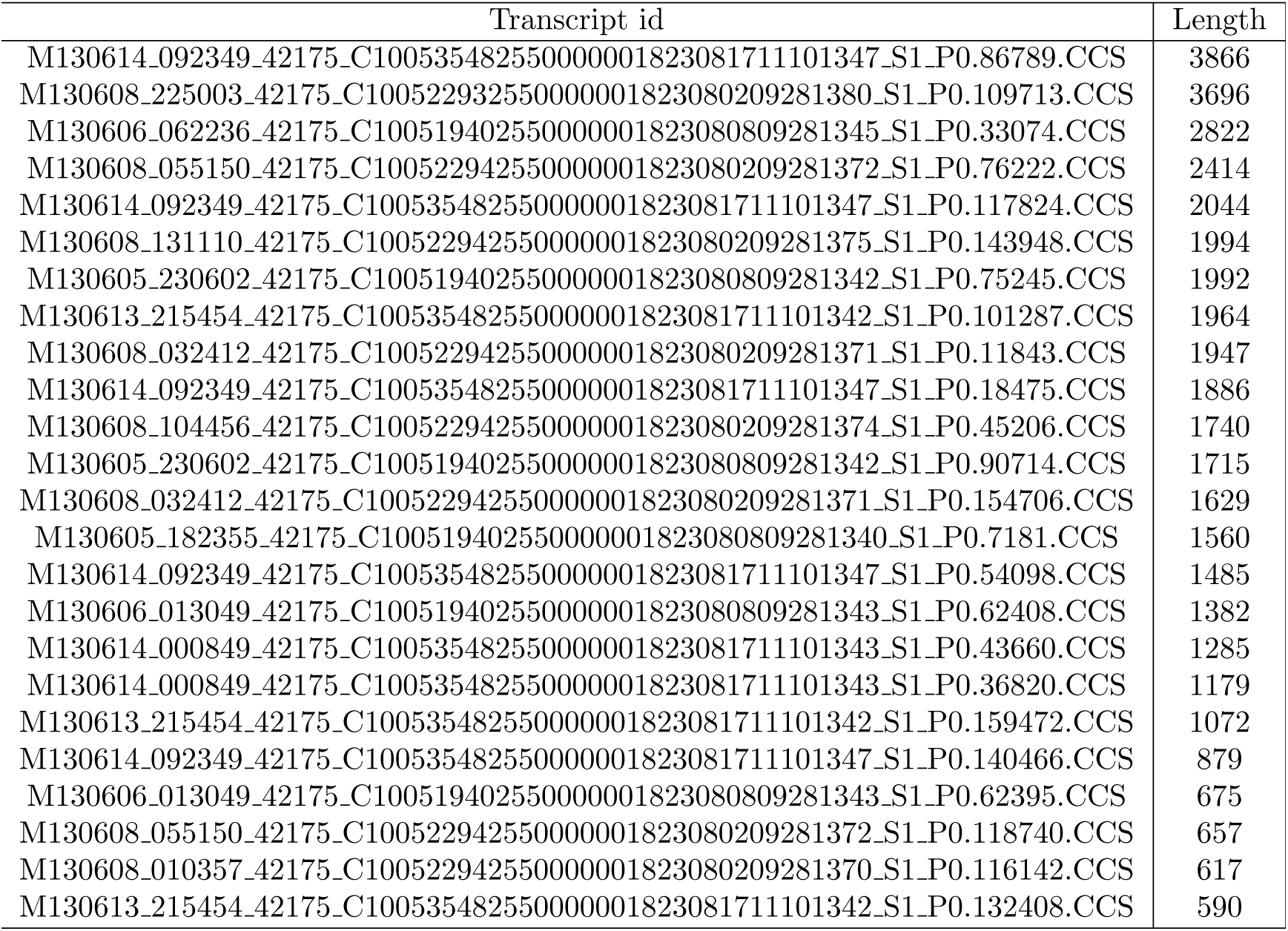
Transcripts (n=24) from GM12878 matching NM_001243.4 (TNFRSF8/CD30) Length of NM_001243.4 = 3706. Summation of counts = 41090. Normalized count = 41090/3706 = ∼11.

### Unannotated genes

Extending previous work identifying un-annotated genes [6], 80 genes were identified which are expressed GM128LCLs, not in hOPTRS, and missing in the GENCODE, RefSeq and RefSeqGene databases (FILE:notannotated.fa). Table 5 shows the annotation of these transcripts obtained from a BLAST on the complete ‘nt’ database.

**Table 5:**
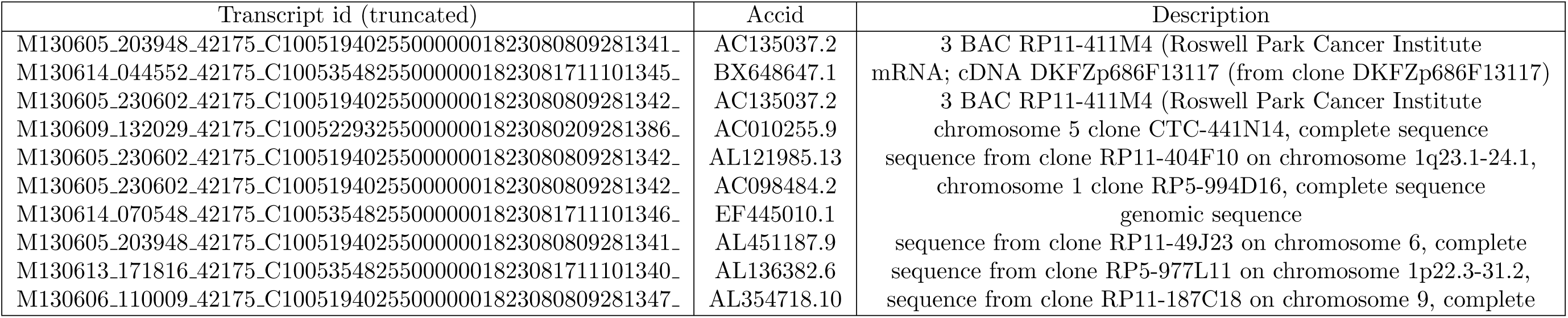
Transcripts having no annotation in GENCODE, RefSeq or RefSeqGene, present in GM128LCLs and absent in hOPTRS. These are the results from doing a BLAST on the complete ‘nt’ database. These transcripts are present in all three LCL cell lines (GM128LCLs), and absent in the transcriptome from twenty tissues (hOPTRS). EValues are 0 for all entries. The complete list (n=82) is in FILE:notannotated.fa.anno.

## Materials and methods

### GENCODE dataset

GENCODE release 25 was obtained from https://www.gencodegenes.org/ (release date 07/2016). Two files - gencode.v25.transcripts.fa (n=200k) and gencode.v25.lncRNA_transcripts.fa (n=27k) - were combined to create a single database (FILE:gencode.v25.ALL.fa.list, n=225785).

### RefSeq dataset

The RefSeq database was created from https://www.ncbi.nlm.nih.gov/nuccore. The current Refseq database has about 200K sequences, and the facet ”genomic DNA/RNA” was ignored (about 20K), leaving ‘facets’ mRNA, rRNA, cRNA, tRNA and ncRNA sequences (FILE:mrna.refseq.180k.fa.list, n=180k). Another set (RefSeqGene) was obtained from ftp://ftp.ncbi.nlm.nih.gov/refseq/H_sapiens/RefSeqGene/(FILE:RefSeqGene.ALL.fa.list, n=6569).

### PacBio transcriptomes

The PacBio transcriptomes from the parent-daughter trio (GM12878, GM12891 and GM12892) were obtained from SRAid:SRP036136 [1]. The Pacbio transcriptome from twenty tissues has been provided at http://www.stanford.edu/∼htilgner/2013_NBT_paper/data/hOP.all.input.ccs.fa.gz [2].

### kmer-based partitioning of ANNODB

GENCODE, RefSeq and RefSeqGene were combined to form a single database (ANNODB), which has redundancies. A kmer-based partitioning algorithm groups the 400K sequences of ANNODB into ∼100k sequences. The clustering algorithm first identifies pairs of sequences having a kmer=100 in common. Finally, a partition is created such that any sequence in a particular cluster has at least one sequence sharing a kmer=100 in the same cluster, and mapping to the same chromosome. The longest sequence in a cluster is chosen as the representative of that cluster. This generic partitioning method can also be done in the case of completely annotated genomes, like RefSeq, by using the gene id.

### kmer-based counts in the transcriptome

Sequences in the transcriptome are kmer=100 matched to the non-partitioned ANNODB. Based on the partitioned ANNODB, counts are generated for the representative sequence. The counts are normalized by summing up the sequence lengths of the transcripts, and dividing it by the length of the representative sequence.

## References

1. Tilgner H, Grubert F, Sharon D, Snyder MP (2014) Defining a personal, allele-specific, and singlemolecule long-read transcriptome. Proceedings of the National Academy of Sciences 111: 9869–9874.

2. Sharon D, Tilgner H, Grubert F, Snyder M (2013) A single-molecule long-read survey of the human transcriptome. Nature biotechnology 31: 1009–1014.

3. Nakatsuka Si, Yutani C, Kurashige M, Kohara M, Nagano T, et al. (2017) An unusual case of epstein-barr virus-positive large b-cell lymphoma lacking various b-cell markers. Diagnostic Pathology 12: 15.

4. Vilain A, Bernardino J, Gerbault-Seureau M, Vogt N, Niveleau A, et al. (2000) Dna methylation and chromosome instability in lymphoblastoid cell lines. Cytogenetic and Genome Research 90: 93–101.

5. Almeida A, Kokalj-Vokac N, Lefrancois D, Viegas-Pequignot E, Jeanpierre M, et al. (1993) Hypomethylation of classical satellite dna and chromosome instability in lymphoblastoid cell lines. Human genetics 91: 538–546.

6. Chakraborty S (2017) Mcf-7 breast cancer cell line pacbio generated transcriptome has ~300 novel transcribed regions, un-annotated in both refseq and gencode, and absent in the liver, heart and brain transcriptomes. bioRxiv: 100974.

7. Reedman BM, Klein G (1973) Cellular localization of an epstein-barr virus (ebv)-associated complement-fixing antigen in producer and non-producer lymphoblastoid cell lines. International Journal of Cancer 11: 499–520.

8. Omi N, Tokuda Y, Ikeda Y, Ueno M, Mori K, et al. (2017) Efficient and reliable establishment of lymphoblastoid cell lines by epstein-barr virus transformation from a limited amount of peripheral blood. Scientific Reports 7.

9. Grassi MA, Rao VR, Chen S, Cao D, Gao X, et al. (2016) Lymphoblastoid cell lines as a tool to study inter-individual differences in the response to glucose. PLoS One 11: e0160504.

10. Sie L, Loong S, Tan E (2009) Utility of lymphoblastoid cell lines. Journal of neuroscience research 87: 1953–1959.

11. Hussain T, Mulherkar R (2012) Lymphoblastoid cell lines: a continuous in vitro source of cells to study carcinogen sensitivity and dna repair. International journal of molecular and cellular medicine 1: 75–87.

12. Kumar S, Curran JE, Glahn DC, Blangero J (2016) Utility of lymphoblastoid cell lines for induced pluripotent stem cell generation. Stem Cells International 2016.

13. Niu N, Wang L (2015) In vitro human cell line models to predict clinical response to anticancer drugs. Pharmacogenomics 16: 273–285.

14. Eid J, Fehr A, Gray J, Luong K, Lyle J, et al. (2009) Real-time dna sequencing from single polymerase molecules. Science 323: 133–138.

15. Quail MA, Smith M, Coupland P, Otto TD, Harris SR, et al. (2012) A tale of three next generation sequencing platforms: comparison of ion torrent, pacific biosciences and illumina miseq sequencers. BMC genomics 13: 1.

16. Nakano K, Shiroma A, Shimoji M, Tamotsu H, Ashimine N, et al. (2017) Advantages of genome sequencing by long-read sequencer using smrt technology in medical area. Human Cell: 1–13.

17. Rhoads A, Au KF (2015) Pacbio sequencing and its applications. Genomics, proteomics & bioinformatics 13: 278–289.

18. English AC, Richards S, Han Y, Wang M, Vee V, et al. (2012) Mind the gap: upgrading genomes with pacific biosciences rs long-read sequencing technology. PloS one 7: e47768.

19. Chakraborty S (2016) Rna-seq assembler artifacts can bias expression counts and differential expression analysis - case study on the chickpea transcriptome emphasizes importance of freely accessible data for reproducibility [version 2; referees: 2 not approved]. F1000Research 5.

20. Steijger T, Abril JF, Engström PG, Kokocinski F, Hubbard TJ, et al. (2013) Assessment of transcript reconstruction methods for rna-seq. Nature methods 10: 1177–1184.

21. Chakraborty S, Britton M, Wegrzyn J, Butterfield T, Martinez-Garcia PJ, et al. (2015). YeATS-a tool suite for analyzing RNA-seq derived transcriptome identifies a highly transcribed putative extensin in heartwood/sapwood transition zone in black walnut.

22. Chakraborty S, Britton M, Martínez-García P, Dandekar AM (2016) Deep RNA-seq profile reveals biodiversity, plant–microbe interactions and a large family of NBS-LRR resistance genes in walnut (juglans regia) tissues. AMB Express 6: 1.

23. Chakraborty S, Martínez-García PJ, Dandekar AM (2016) Yeatsam analysis of the walnut and chickpea transcriptome reveals key genes undetected by current annotation tools. F1000Research 5.

24. Martínez-García PJ, Crepeau MW, Puiu D, Gonzalez-Ibeas D, Whalen J, et al. (2016) The walnut (juglans regia) genome sequence reveals diversity in genes coding for the biosynthesis of nonstructural polyphenols. The Plant Journal .

25. Lagarde J, Uszczynska-Ratajczak B, Santoyo-Lopez J, Gonzalez JM, Tapanari E, et al. (2016) Extension of human lncrna transcripts by race coupled with long-read high-throughput sequencing (race-seq). Nature communications 7.

26. Skalsky RL (2017) Analysis of viral and cellular micrornas in ebv-infected cells. Epstein Barr Virus: Methods and Protocols: 133–146.

27. Teng G, Papavasiliou FN (2009) Shhh! silencing by microrna-155. Philosophical Transactions of the Royal Society of London B: Biological Sciences 364: 631–637.

28. Gatto G, Rossi A, Rossi D, Kroening S, Bonatti S, et al. (2008) Epstein-barr virus latent membrane protein 1 trans-activates mir-155 transcription through the nf-*κ*b pathway. Nucleic acids research 36: 6608–6619.

29. Yin Q, McBride J, Fewell C, Lacey M, Wang X, et al. (2008) Microrna-155 is an epstein-barr virus-induced gene that modulates epstein-barr virus-regulated gene expression pathways. Journal of virology 82: 5295–5306.

30. Eis PS, Tam W, Sun L, Chadburn A, Li Z, et al. (2005) Accumulation of mir-155 and bic rna in human b cell lymphomas. Proceedings of the National Academy of Sciences of the United States of America 102: 3627–3632.

31. Tobinai K, Klein C, Oya N, Fingerle-Rowson G (2016) A review of obinutuzumab (ga101), a novel type ii anti-cd20 monoclonal antibody, for the treatment of patients with b-cell malignancies. Advances in Therapy: 1–33.

32. Solano-Genesta M, Tarín-Arzaga L, Velasco-Ruiz I, Lutz-Presno JA, González-Llano O, et al. (2012) Cd20 expression in b-cell precursor acute lymphoblastic leukemia is common in mexican patients and lacks a prognostic value. Hematology 17: 66–70.

33. Enukashvily N, Donev R, Waisertreiger IR, Podgornaya O (2007) Human chromosome 1 satellite 3 dna is decondensed, demethylated and transcribed in senescent cells and in a431 epithelial carcinoma cells. Cytogenetic and genome research 118: 42–54.

34. Qu Gz, Grundy PE, Narayan A, Ehrlich M (1999) Frequent hypomethylation in wilms tumors of pericentromeric dna in chromosomes 1 and 16. Cancer genetics and cytogenetics 109: 34–39.

35. Orsetti B, Nugoli M, Cervera N, Lasorsa L, Chuchana P, et al. (2006) Genetic profiling of chromosome 1 in breast cancer: mapping of regions of gains and losses and identification of candidate genes on 1q. British journal of cancer 95: 1439–1447.

36. Frankish A, Uszczynska B, Ritchie GR, Gonzalez JM, Pervouchine D, et al. (2015) Comparison of gencode and refseq gene annotation and the impact of reference geneset on variant effect prediction. BMC genomics 16: S2.

37. Thorley-Lawson DA, Schooley RT, Bhan AK, Nadler LM (1982) Epstein-barr virus superinduces a new human b cell differentiation antigen (b-last 1) expressed on transformed lymphoblasts. Cell 30: 415–425.

38. Hu S, Xu-Monette ZY, Balasubramanyam A, Manyam GC, Visco C, et al. (2013) Cd30 expression defines a novel subgroup of diffuse large b-cell lymphoma with favorable prognosis and distinct gene expression signature: a report from the international dlbcl rituximab-chop consortium program study. Blood 121: 2715–2724.

